# Barnacle encrustation on molluscan prey offers associational resistance against drilling predation

**DOI:** 10.1101/2020.06.08.139931

**Authors:** Vigneshbabu Thangarathinam, Devapriya Chattopadhyay

## Abstract

Predation is one of the driving forces that shaped the marine ecosystems through time. Apart from the anti-predatory strategies adopted by the prey, the predatory outcome is often indirectly influenced by the other members of the ecological community. Association between organisms are often found to influence the outcome and the evolution of such association may have been guided by such interactions. Mollusc-burnacle association, although common, is not explored to assess if the epibiont offers the molluscs any protection against predation (associational resistance) or increases the risk by attracting predators (shared doom). Using a series of controlled experiments with a drilling predator (*Paratectonatica tigrina*), its prey (*Pirenella cingulata*) and an epibiont (*Amphibalanus amphitrite*), we evaluated the effect of epibionts on the drilling behavior of the predator by documenting the successful attack (Drilling frequency, DF), and handling time. Our results show that the prey with epibionts are significantly less likely to be drilled when the predator has sufficient choice of prey, consistent with the tenets of the associational resistance. The preference of choosing the non-encrusted prey, however, diminishes with fewer available prey. The handling time is significantly higher in the attacks on the encrusted prey than non-encrusted prey, even though the barnacles are not drilled. Although the proximity of the drilling site to encrustation tends to increase the handling time, the size of encrustation does not have any effect. Because the profitability of prey largely depends on the ratio of handling time and the energetic yield from consuming the prey, the increase in handling time due to encrustation makes it less profitable for the predator. The role of encrustation as a deterrent to predation might also explain the complex shell architecture in some prey gastropods that increases the likelihood of encrustation besides providing direct resistance against predation.

## Introduction

Organisms interact with each other in various direct and indirect ways, such as predation, parasitism, symbiosis, and competition. Complexity and diversity in the interactions between organisms are suggested to be the primary driving force behind the structure of ecological communities and food webs [1]. Predation is one of the critical biotic interactions that significantly influenced the marine ecosystems in deep time [2, 3], and hence, the predatory marks that survive in the fossil record is of great interest to paleontologists. Fossil records of predation in marine organisms is dominated by the occurrence of predatory drill holes created by carnivorous gastropods [3]. The success of drilling predation often depends on the anti-predatory strategies adopted by the prey including morphological modification (such as producing armored shells, developing toxins) and behavioral changes (such as clumping, mimicking other dangerous prey) [3]. Apart from the anti-predatory traits of the prey, predatory success is also influenced by other biotic interactions. It has been shown that predatory behavior of a drilling gastropod is affected by the presence of a secondary predator [4] and intraspecific competition [5, 6, 7, 8]. Yet another biotic association, epibiosis may potentially affect predation where the epibiont lives attached to the exterior of another organism. Epibiosis on molluscan shells are known to conceal cemented prey from predators [9, 10, 11] and deter durophagous predators [12]. In some cases, epibionts are also harmful to the molluscan host by reducing their mobility and reproductive rate [13].

In spite of the general interest in drilling predation because of its deep-time quantifiable record, the influence of epibiosis on drilling predation has not been explored. Using a series of controlled experiments with a drilling predator (*Paratectonatica tigrina*), its natural prey (*Pirenella cingulata*) and a common epibiont (*Amphibalanus amphitrite*), we evaluated if the epibiont offers the molluscan prey any protection against drilling predation (associational resistance) or increases the risk by attracting predators (shared doom).

## Materials and method

### Specimen collection

A total of 300 live gastropod specimens representing two gastropod species (*Paratectonatica tigrina* and *Pirenella cingulata*) (Fig. 1A-C) were collected from the tidal flats of Chandipur, Odisha from two spots (N21°27’19.5”, E87°02’55.5” and N21°26’43.8”, E87°02’56.7”) during three seasons of fieldwork (Summer 2019, 27-29^th^ May 2019, 21^st^ September 2019). An established alien barnacle epibiont, *Amphibalanus amphitrite* is often associated with these shells [14]. Majority of the encrustations were alive when collected. For a few of them, we removed the soft tissue of the encrustation using a pair of metal tweezers and considered them as dead encrustation.

**Fig. 1.**
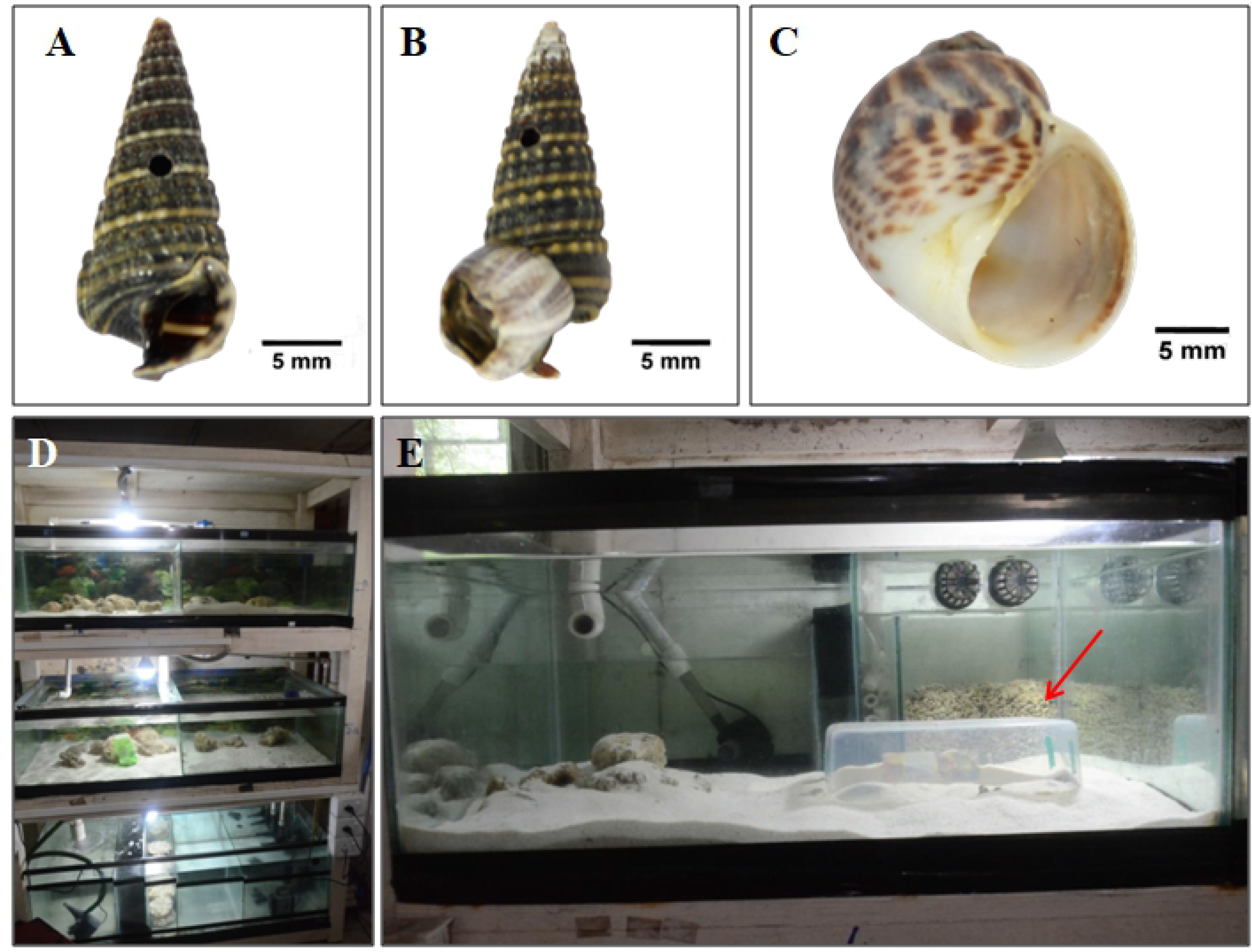
Specimens used in the experiment including the prey species (*Pirenella cingulata*) without encrustation (A), with encrustation (B), and predatory gastropod *Paratectonatica tigrina* (C). Experimental setup with a circulating tanks (D) and the experiment cage (marked by an arrow) inside a tank (E).

### Experimental setup

For this experiment, we used the synthetic saltwater aquarium housed in Ecological Field Station, IISER Kolkata (Fig 1D). The physical conditions were kept constant throughout the experiment (with the temperature at 25°C, the salinity at 33.4 psu, and the pH at 8.4). The specimens were all initially kept at the recirculation tank for ten days for acclimatization and later transferred to different chambers within the recirculating tank during the experiment. The dimensions of the experimental chambers were 1.2 m × 0.5 m × 0.5 m.

### Experimental design

One individual of *Paratectonatica tigrina* was kept in an underwater plastic cage (204mm x 118mm x 83mm) (Fig. 1E) for one day. The cuboid cage had three holes on each side to facilitate water circulation. Different combinations of prey specimens were introduced to the cage, depending on the type of the experiment (Table 1). The experimental setup was monitored once every six hours. Dead prey individuals were also removed at the same interval. The removed specimens were not replaced. We continued each experiment for five days. We documented the size of the involved specimens, the order of the drilling predation, the time taken for each predation, the size and the position of drill holes, the location of the encrustation.

**Table 1.** Experimental design.

| Experiment | Aim | Trials | Trials with occurrence of predation | Involved groups |  |
| --- | --- | --- | --- | --- | --- |
|  |  |  |  | Predator | Prey |
| 1 | Comparison of drilling on prey with and without encrustation | 27 | 23 | *N. tigrina* = 27 | *P. cingulata* with live encrustation =69, *P. cingulata* without encrustation =69 |
| 2 | Comparison of drilling on prey with live and dead encrustation | 10 | 10 | *N. tigrina* = 10 | *P. cingulata* with live encrustation =30, *P. cingulata* with dead encrustation =30 |

All the prey gastropods from the trials in which no predation occurred were reused in the subsequent trials.

#### Experiment 1: Effect of the presence of encrustation

Then three individuals of *Pirenella* with live encrusters and three individuals of *Pirenella* without encrusters were introduced into the cage containing the predator. In every observation, the predation by the *Paratectonatica* on the *Pirenella* was noted.

#### Experiment 2: Effect of the nature of the encrustation

Three prey specimens with dead barnacles on their shell, and the other three prey specimens with live barnacles were introduced to the cage containing the predator. Observations were made at 6-hour intervals for five days.

### Analysis

Prey gastropod (*Pirenella*) shells with complete drill holes are considered as drilled specimens. Barnacles with no operculum or soft tissue were considered dead. Drilling frequency is calculated by dividing the number of drilled specimens by the total number of prey gastropods. Handling time was considered as the time duration for which the predator was found attached to the prey. Handling time was not distinguished into drilling time and consumption time [15] because the predation was not interrupted during observations. The degree of encrustation was calculated as the ratio of the diameter of the encrustation to the length of the host gastropod.

Two-tailed Chi-square test was done on the drilling frequencies across different categories of prey. The Welch’s t-test were done to compare sizes and handling time between prey of different categories. All statistical tests were performed in R with tidyverse package [16,17] and XLSTAT [18].

## Results

Out of the total of 198 *Pirenella cingulata* prey, 129 were encrusted. Out of the encrusted *Pirenella cingulata* prey, 59 were drilled (Table 2). All drill holes were complete. Except for one, none of the drillings were directly on the encrustation. None of the gastropods died due to causes other than drilling predation during these experiments.

**Table 2.** Outcome of the drilling experiments. Results with statistical significance is marked bold.

| Experiment | Encrustation | First attack |  | Chi-square test | Subsequent attacks |  | Chi-square test |
| --- | --- | --- | --- | --- | --- | --- | --- |
|  |  | Drilled | Undrilled |  | Drilled | Undrilled |  |
| 1 | Encrusted (live) | 6 | 17 | $\chi^2 = 10.5217$,<br>**p = 0.001** | 30 | 16 | $\chi^2 = 0.415$, p =<br>0.519 |
|  | Non-encrusted | 17 | 6 |  | 27 | 19 |  |
| 2 | Encrusted (live) | 6 | 4 | $\chi^2 = 0.8$, p =<br>0.371 | 7 | 13 | $\chi^2 = 0.114$, p<br>=0.736 |
|  | Encrusted (dead) | 4 | 6 |  | 6 | 14 |  |

In the experiment comparing prey with encrusters (live) and prey without encrusters, 27 trials were carried out. In the 23 trials, 80 out of 138 prey gastropods were drilled (DF = 0.58, Table 2). The encrusted prey had significantly lower DF (DF = 0.52) than that of the non-encrusted prey (DF = 0.64, Table 2) for the 1^st^ attacks (Chi-square test, χ^2^ = 10.5217, p = 0.001, Table 2, Fig. 2A). All the subsequent predations (with less than five prey), however, do not show such significant difference (Table 2, Fig. 2A). The drilling frequency does not change significantly (χ^2^ = 0.4551. p = 0.929) for prey with different amount of encrustation (Table 2, Fig. 2B). In four out of the 27 trials, no predation happened within the five-day experimental period.

**Fig. 2.**
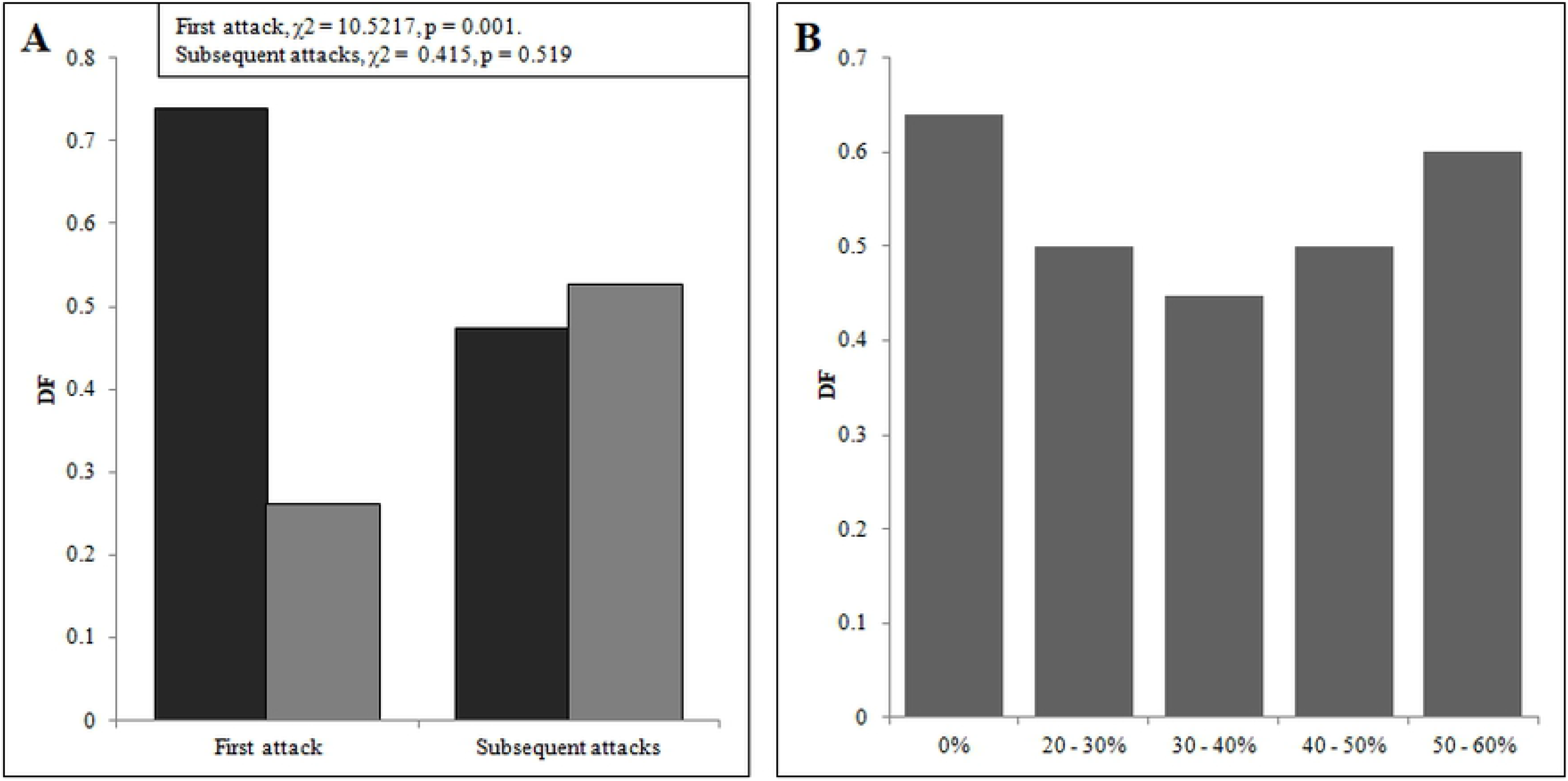
Plot showing the difference in the DF between prey with and without encrustation in different events of predation (A) and with different extent of encrustation (B). Black bars represent prey without encrustation and dark grey bars represent prey with encrustation.

The comparison between the handling time of encrusted and non-encrusted prey shows a significant difference where the average handling time is longer for encrusted prey (Welch’s t-test, t = 2.0993, p = 0.04) (Fig. 3A). This difference cannot be explained by the size of the prey because the size distribution of the prey with and without encrustation is indistinguishable (Welch’s t-test, t = 0.125, p = 0.9) (Fig. 3B, Table 3).

**Fig. 3.**
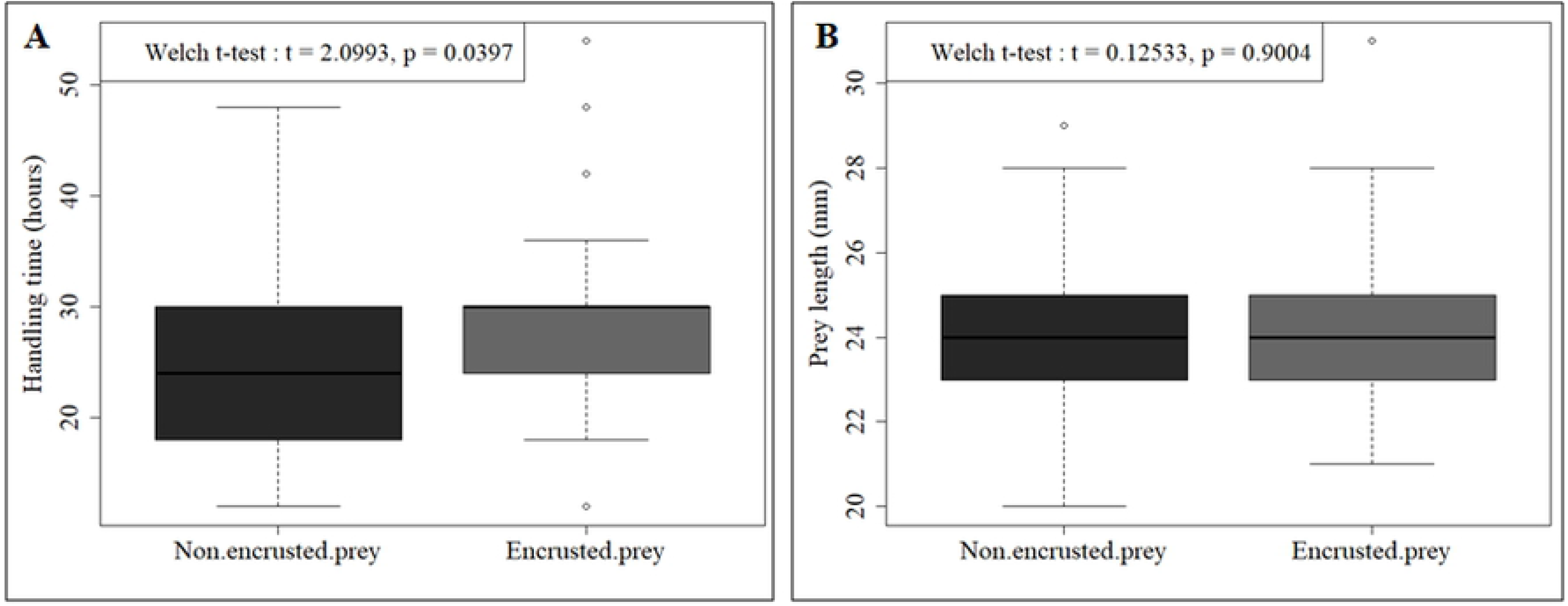
Box plots comparing handling time (A) and prey length (B) of non-encrusted and encrusted prey. The boxes are defined by 25th and 75th quantiles; thick line represents the median value. Light grey bars represent prey with dead encrustation and dark grey bars represent prey with live encrustation.

**Table 3.**
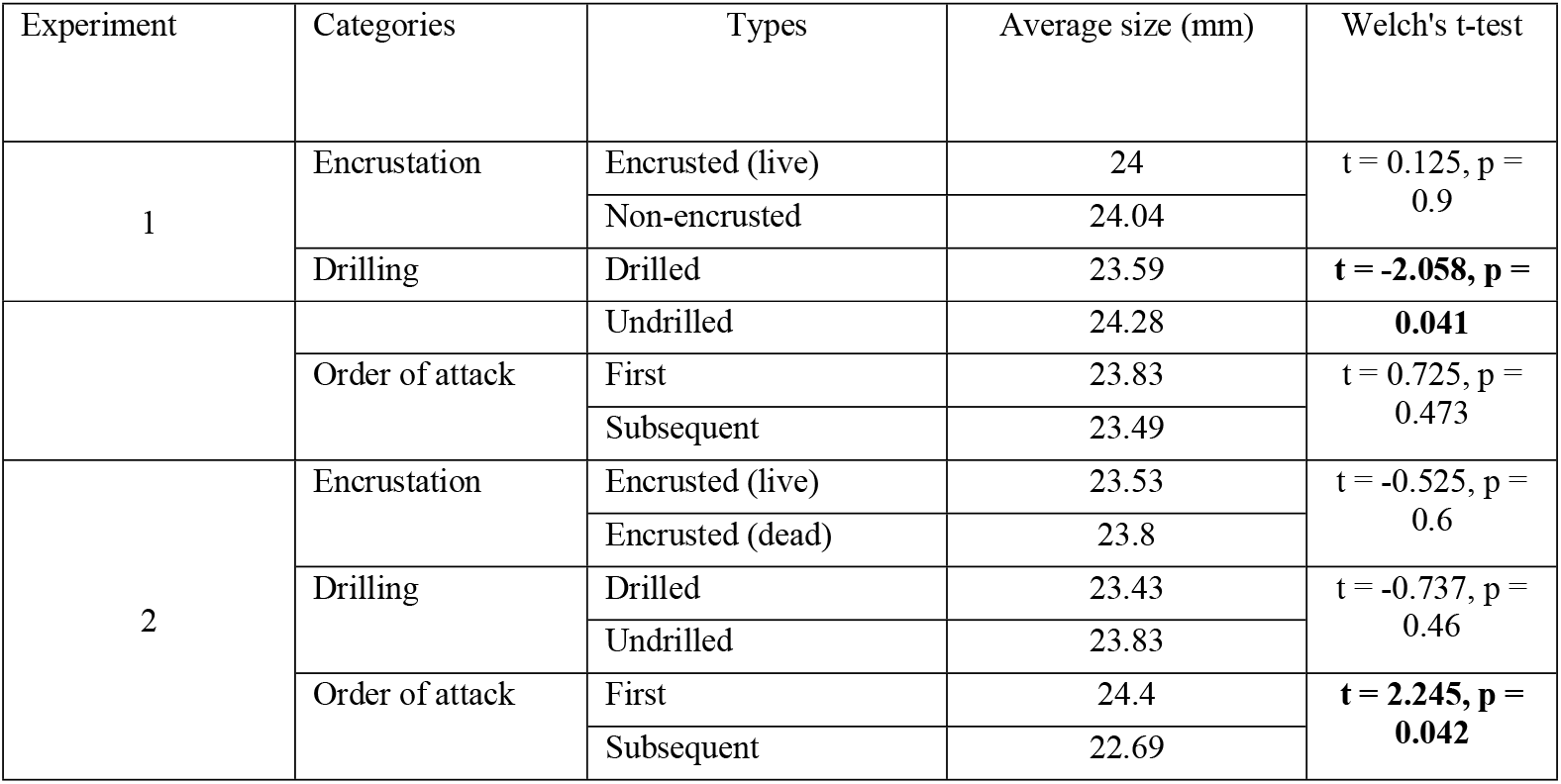
A comparison of size between different categories of prey used in the experiment. Results with statistical significance is marked bold.

In the experiments comparing prey with dead encrusters and prey with live encrusters, ten trials were done. Predation was observed in all ten trials and a total of 23 out of 60 prey gastropods were drilled (DF=0.38, Table 2, Fig. 4A). The drilling frequency for prey with live encrustation is not significantly different from that of the dead encrustation in the 1st attack and all the subsequent ones (Chi-square test, χ2 = 0.282, p = 0.595) (Fig. 4A, Table 2).

**Fig. 4.**
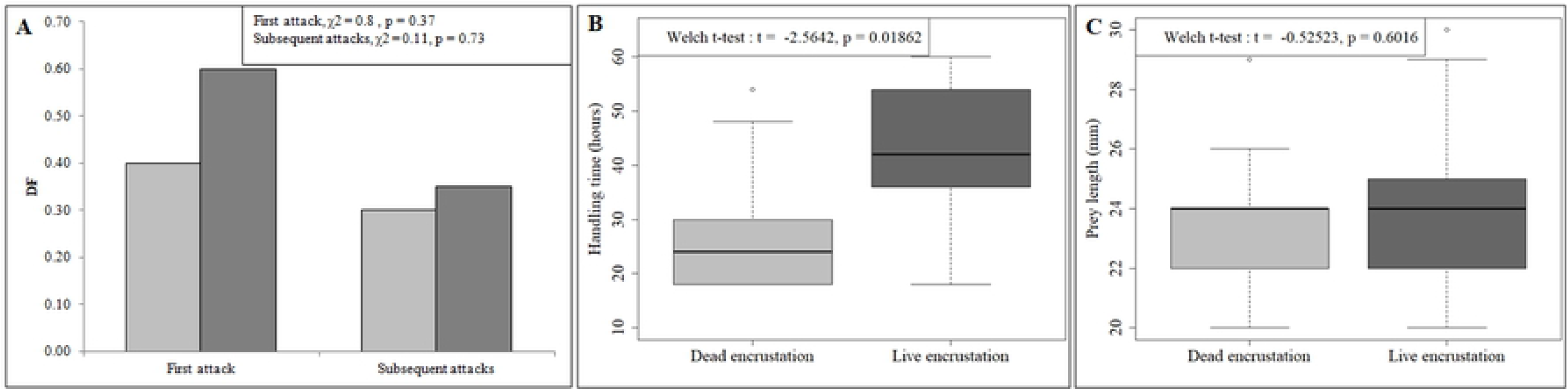
Plot showing the difference in the DF between prey with dead encrustation and prey with live encrustation in different events of predation (A). Box plots comparing the predation events with live and dead encrustation in terms of the prey length (B) and the handling time (C). The boxes are defined by 25th and 75th quantiles; thick line represents the median value. Light grey bars represent prey with dead encrustation and dark grey bars represent prey with live encrustation.

The comparison between handling time for prey with live and dead encrustation shows a significant difference with significantly longer average handling time for prey with live barnacles (t = −2.5642, p = 0.02) (Fig.4B). This difference cannot be explained by the size of the prey because the size distribution of the prey with live and dead encrustation is statistically indistinguishable (Welch’s t-test, t = −0.525, p = 0.6) (Table 3, Fig.4C).

The handling time shows a negative correlation with the distance of the drillhole from the encrustation for all drilled prey with attachment (Fig. 5A). The handling time, however, does not show any correlation with the size of the encrustation (Fig. 5B).

**Fig. 5.**
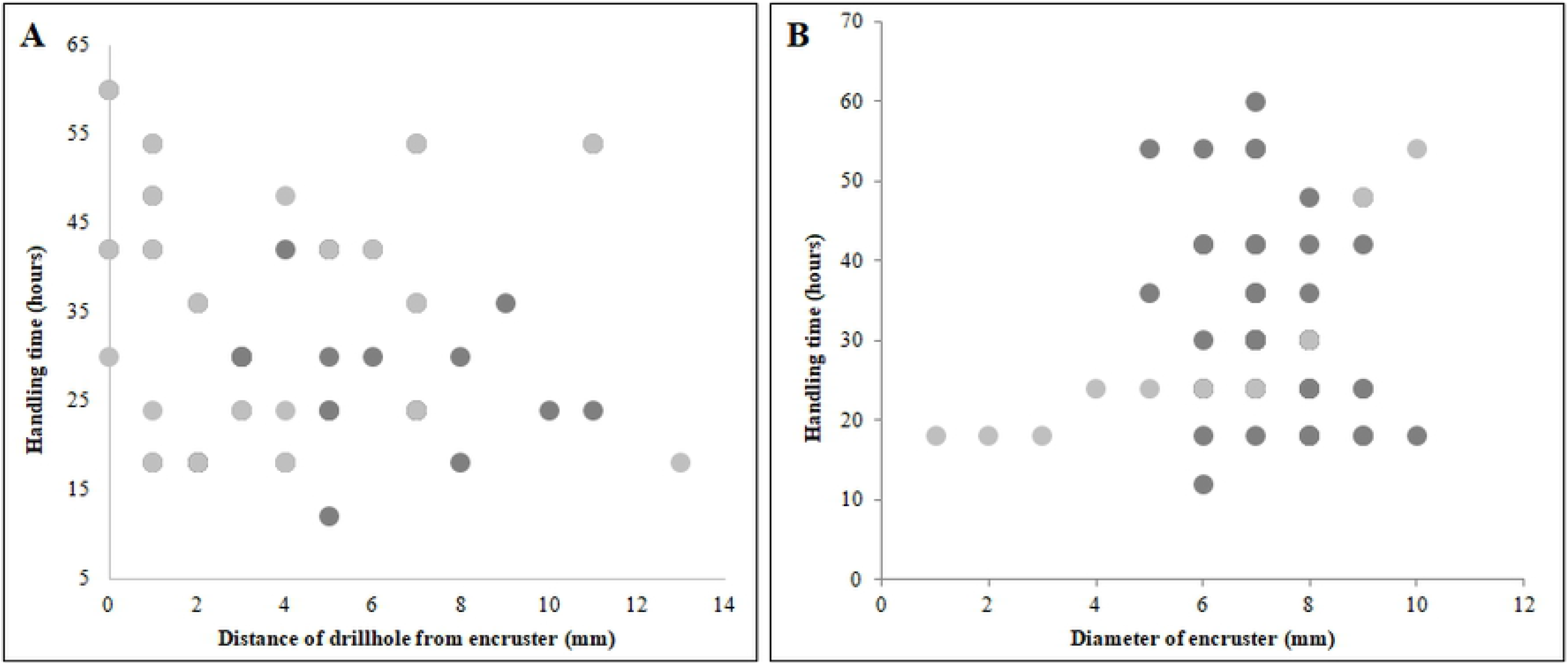
Plots showing the relationship of handling time with the distance of the drill hole from the encruster (A), and the diameter of encruster (B). Light grey dots and light grey line represent prey with dead encrusters and dark grey dots and dark grey line represent prey with live encrusters.

## Discussion

### Associational resistance or shared doom

Susceptibility of a species to predation does not always depend only on its profitability as food and anti-predatory defenses. More often than not, the predatory behavior depends on the community aspects of an assemblage (such as the trophic structure, presence of other predators) [4, 19]. The susceptibility of the prey species is, therefore, determined by its properties (such as anti-predatory defense and profitability) relative to the characteristics of other members of the community [20]. There are two commonly predicted outcome that a prey species can experience through an association: a) positive (associational resistance scenario) [21, 22, 23] and b) negative (shared doom scenario) [20]. An associational resistance describes a scenario where a preferred species may experience reduced risk because of its association with more preferred species or proximity to an unpalatable (even dangerous) species. The shared doom scenario, in contrast, portrays a situation where the risk of the prey increases due to its association with a species that attracts the predator. Although these scenarios were primarily designed to model herbivory [24, 25], it can be used to understand the effect of epibiosis on the predation of the marine shelled prey. The effect of barnacle epibionts reducing the success of a drilling predator, as shown in the present study, supports the associational resistance, where an epibiont reduces the chance of predation on its host, by changing its host’s exterior such that it is not consumed as easily by the predator [26, 27]. Apart from observing an effective associational resistance against drilling predation, our experiments also point to the dependence of the predatory behavior on the order of the attack; the predator showed a significant preference towards non-encrusted prey in the first predation of each trial (Fig. 2, Table 2). The two factors changing with the order of the predation, level of hunger of the predator and number of available prey, have the potential to influence the predatory behavior. It is common for drilling predator to change their behavior with hunger, especially in prey preference [28, 29, 30]. All of these studies, however, found a less selective behavior of the predator with increased hunger. The drop in prey-preference with progressively satiated predator in our experiment is at odds with the expectation. The availability of large number of prey often shows the true behavioral traits of the predators that are hard to observe when the prey choice is limited. Enderlein et al [31] in their study on the effect of epibionts on crab predation, only took observation of the 1^st^ attack and stopped the experiment immediately (p. 236) probably to avoid such an issue. The lack of selective predation in subsequent attacks is probably due to the insufficient prey choice.

### The relative contribution of the predator, prey and the epibiont

The relative importance of each member (predator, prey, and the epibiont) in deciding the outcome of this tripartite game is not well understood. Prey type tends to have a limited role in modulating the risk (increase or decrease) by association. Shelled invertebrates demonstrated both associational resistance and shared doom mainly depending on the type of the predator and their mode of detection (Table 4, Fig. 6). Unlike the prey, the predator plays an important role in deciding the outcome of predation in the marine ecosystem. There are three major predatory groups for shallow benthic prey, namely, the durophagous predators (such as crabs, lobsters, fishes, etc.), sea stars, and gastropods. The durophagous predators primarily detect prey by visual and chemical cues, and their predation success seems to be affected by the presence of epibionts. Majority of cases of such association support associational resistance where the durophagous predators avoid the prey or show a reduced success. A variety of crabs demonstrated such avoidance [12, 31, 32, 33]. The crab consumes an encrusted prey in some cases, always after removing the epibiont, indicating the negative effect of the chemical cues on the durophagous predator produced by the epibionts [12]. Lobsters are also known to preferentially attack shelled prey without encrustation primarily because the epibionts act as an effective camouflage that conceals the prey [9, 34]. Fishes (rays and sharks) are also known to avoid prey with encrustation [9]. Predatory behavior of a variety of echinoderms (sea star, urchin) also supports the associational resistance [11]. Only on rare occasions, such predation is found not to be affected by the presence of epibionts [35]. Predation by non-drilling gastropods is also found to support associational resistance [9]. The behavior of drilling predator, however, has not been explicitly tested before. Our results demonstrate a predatory behavior that shows signs of associational resistance.

**Table 4.** A compilation of the recorded effects of epibionts on predation events.

| Predation type | Predatory species | Predator type | Prey species | Epibiont type | Outcome | Reference |
| --- | --- | --- | --- | --- | --- | --- |
| Drilling | *Natica tigrina* | Mollusca (Gastropoda) | *Cerithidae cingulata* | Barnacle | Associational resistance | Present study |
| Crushing | *Carcinus maenas* | Arthropoda (Decapoda) | *Mytilus edulis* | Barnacle | Shared doom | Enderlein et al. 2003 |
|  | *Carcinus maenas* | Arthropoda (Decapoda) | *Littorina littorea* | Barnacle | Shared doom | Buschbaum and Reise, 1999 |
|  | *Carcinus maenas* | Arthropoda (Decapoda) | *Mytilus edulis* | Barnacle | Shared doom | Wahl et al 1997 |
|  | *Calappa flammea* | Arthropoda (Decapoda) | *Pagurus pollicaris* | Coral | Associational resistance | McLean and Mariscal 1973 |
|  | *Carcinus maenas* | Arthropoda (Decapoda) | *Mytilus edulis* | Hydrozoa | Associational resistance | Wahl et al 1997 |
|  | *Panulirus argus* | Arthropoda (Decapoda) | *Spondylus americanu* | Sponges | Associational resistance | Feifarek 1987 |
|  | *Carcinus maenas* | Arthropoda (Decapoda) | *Mytilus edulis* | Tunicate worm | Associational resistance | Aucker et al. 2014 |
|  | *Diodon hystrix* | Chordata (Actinopterygii) | *Spondylus americanu* | Sponges | Associational resistance | Feifarek 1987 |
|  | *Aetobatis narinari* | Chordata (Elasmobranchii) | *Spondylus americanu* | Sponges | Associational resistance | Feifarek 1987 |
|  | *Dasyatis americanus* | Chordata (Elasmobranchii) | *Spondylus americanu* | Sponges | Associational resistance | Feifarek 1987 |
| Ingestion | *Pycnopodia helianthoides* | Echinodermata (Asteroidea) | *Chlamys hastata* | Barnacle | Shared doom | Donovan et al. 2003 |
|  | *Pycnopodia helianthoides* | Echinodermata (Asteroidea) | *Chlamys hastata* | Barnacle | Shared doom | Farren and Donovan, 2007 |
|  | *Asterias rubens* | Echinodermata (Asteroidea) | *Mytilus edulis* | Barnacle | No significant effect | Caderwood et al. 2015 |
|  | *Asterias rubens* | Echinodermata (Asteroidea) | *Mytilus edulis* | Barnacle | Associational resistance | Laudien and Wahl 2004 |
|  | *Asterias rubens* | Echinodermata (Asteroidea) | *Mytilus edulis* | Hydrozoa | Associational resistance | Laudien and Wahl 2004 |
|  | *Asterias rubens* | Echinodermata (Asteroidea) | *Mytilus edulis* | Sponges | Associational resistance | Laudien and Wahl 2004 |
|  | *Pycnopodia helianthoides* | Echinodermata (Asteroidea) | *Chlamys hastata* | Sponges | Associational resistance | Farren and Donovan, 2007 |
|  | *Pycnopodia helianthoides* | Echinodermata (Asteroidea) | *Chlamys hastata* | Sponges | Associational resistance | Bloom 1975 |
|  | *Orthasterias koehleri* | Echinodermata (Asteroidea) | *Chlamys hastata* | Sponges | Associational resistance | Bloom 1975 |
|  | *Pycnopodia helianthoides* | Echinodermata (Asteroidea) | *Chlamys hastata* | Sponges | Associational resistance | Farren and Donovan, 2007 |
|  | *Pisaster* | Echinodermata | *Chlamys* | Sponges | Associational | Bloom 1975 |
|  | *ochraceus* | (Echinoidea) | *hastata* |  | resistance |  |
| Peeling | *Fasciolaria tulipa* | Mollusca (Gastropoda) | *Spondylus americanu* | Sponges | Associational resistance | Feifarek 1987 |

**Fig. 6.**
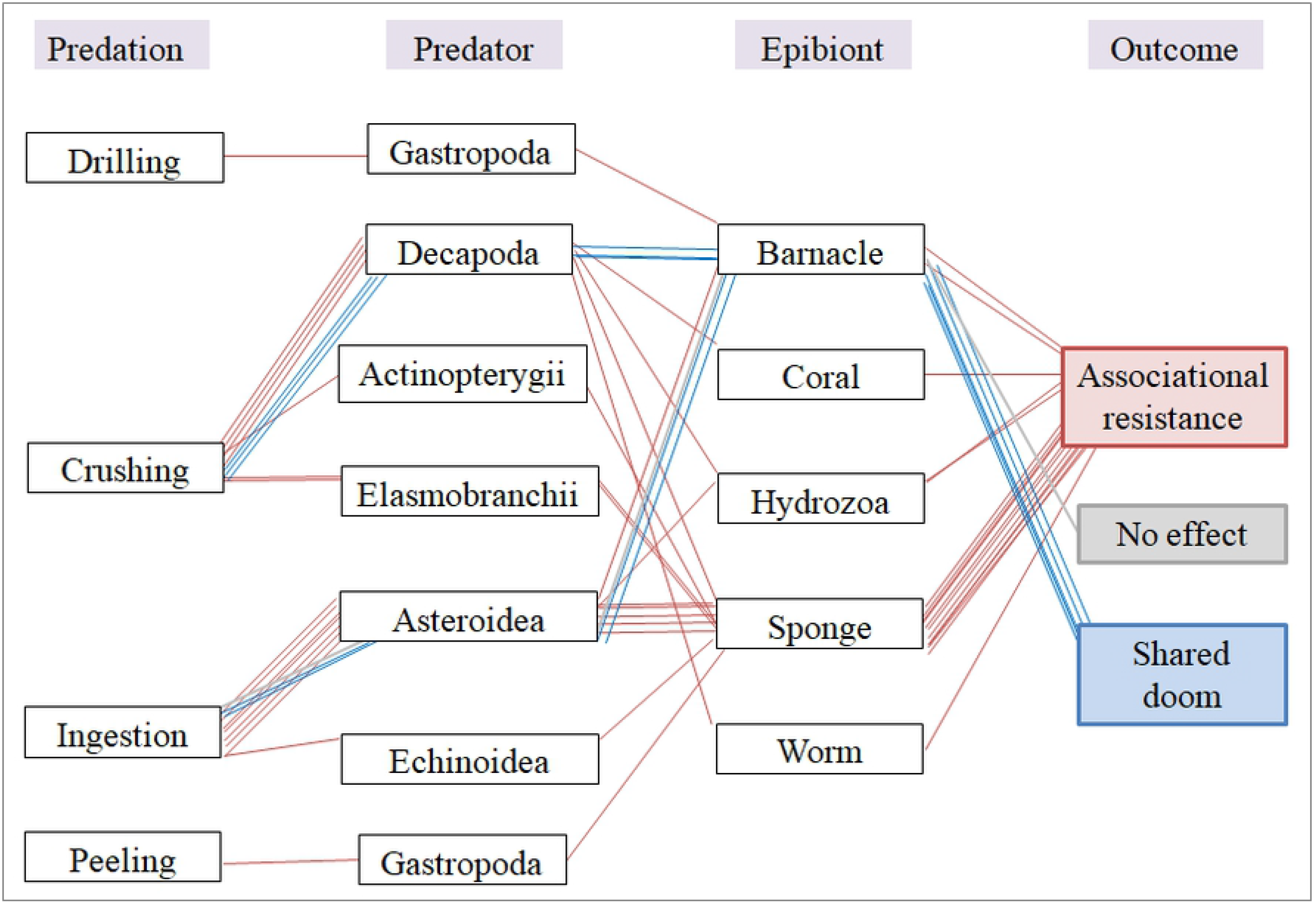
Diagram showing the two outcomes of from various associations of predator and epibiont based on the published literature and the present study. The red and blue lines represent the interactions that resulted in “Associational resistance” and “Shared doom” respectively. Grey line represents no effect.

Type of the epibiont often plays an essential role in modulating predatory behavior. Some epibionts offer protection by increasing shell thickness (such as bryozoan, barnacle) and offer effective visual camouflage [9, 34] while others offer tactile camouflage (such as anthozoan, hydrozoan, tunicate) [12, 31, 32, 33] aiding the associational resistance. The results of our study indicate that the barnacle can also offer tactile camouflage [12] afflicting the drilling gastropods with the limited vision that primarily detects the prey by chemical cues. Although we did not find any difference in prey-preference between prey with live and dead encrustation, such lack of preference can be explained by the way dead encrustation were created. Because the dead encrustations in our experiment were freshly dead and there could have been residual soft tissue emanating the chemical cues, it might be difficult for the drilling predator to distinguish between live and dead encrustation.

Barnacles are also known to directly affect their hosts by decreasing mobility and hindering copulation [36]. Such a reduction in mobility may lead to an increase in predation risk. Apart from directly reducing the fitness of their host, barnacles can increase predation risk for the host they settle on by their mere presence. Enderlein et al. [31] found that the crab *Carcinus maenas* prefers barnacle-encrusted *Mytilus edulis* mussels over clean mussels because the barnacles make the prey more comfortable to handle.

Although there is no direct observation to assess the effect of barnacle association on drilling predation in ecological settings, indirect evidences tend to support associational resistance. A study on the drilling behavior of Naticidae gastropods from the intertidal zone of Chandipur, India showed incidence of naticid drilling on barnacles in association with bivalve *Timoclea imbricata* [37]. Although the barnacles were drilled, we can compute the attack frequency on such a bivalve-barnacle association. By comparing the published data on naticid predation on *Timoclea* without encrustation from the same intertidal habitat of Chandipur we found that the bivalve-barnacle association has a lower probability of getting drilled (DF=0.23, 37) in comparison to non-encrusted prey (DF=0.35, 8) and this difference is statistically significant (Chi-square test, p=0.04) (Table 5). This ecological data also implies an associational resistance where the presence of a barnacle encruster reduces the chances of drilling predation for the molluscan host. It is interesting to note that barnacle themselves can experience “shared doom”. The Chandipur intertidal assemblage shows a relatively higher incidence of drilling on encrusted barnacles when they are in association with *Timoclea* [37]. Naticids of that ecosystem was rarely found to attack the barnacles without such association [38]. This showcases a “shared doom” scenario for barnacle prey where their association with the molluscs increases their risk from predation.

**Table 5.** A comparison of the predation record on bivalve *T. imbricata* from Chandipur. The drilling record of Mondal et al (2019) is on attached barnacles.

|  | Attacked individuals | Unaffected individuals | DF | Reference |
| --- | --- | --- | --- | --- |
| With encrustation | 303 | 1007 | 0.23 | Mondal et al, 2019 |
| Without encrustation | 92 | 217 | 0.35 | Chattopadhyay et al, 2014 |

In the context of the most recent climate change predictions, such interactions become more relevant [31]. For the temperate regions an increase in temperature and eutrophication, a decrease in salinity and an enhanced invasiveness by warm-adapted species are expected (IPCC, 2001a, b). In one hand, less saline conditions will reduce shell thickness in molluscs increasing the predation risk, it will also facilitate the arrival of new epibiont species and may decrease predation risk by associational resistance. This may lead to unpredictable patterns of predator-prey interactions.

### Mechanism of associational resistance

The mechanism of an epibiont in dissuading a predator was primarily thought to be by offering visual camouflage or changing the effluence profile of the prey/host. Our study identifies yet another aspect – the handling time. We found that the presence of barnacles, especially the live ones, significantly increases the handling time (Fig. 3). Previous studies on barnacle predation often found a low incidence of naticid drilling [39]. Naticids typically envelop their prey in mucus and orient them preferentially [40]. Such handling was thought to be impossible to execute with cemented barnacles like the cemented bivalves [41]. Lack of prey manipulability was, therefore, considered as the causal mechanism for the apparent lack of naticid drilling on barnacles [39] without experimental verification. Studies on barnacles settled on mobile prey, however, shows a higher incidence of naticid drilling. Our study reveals that presence of barnacle encrustation significantly increases the handling time for drilling predation. Handling time plays an important role in the energy maximization model developed from the optimal foraging theory [42]; it is a measure of the cost incurred during a predation and its increase implies a decrease in the relative profitability of a specific prey to a predator [43]. The increase in handling time can be due to the increased recognition time or increase in time taken by the predators to find a suitable drilling site. We do not find any evidence for encrustation increasing ease of prey handing unlike the previous claims [31] where predatory crabs are found to prefer mussels with encrustation (even with synthetic mimics of the encrustation). Unlike the crabs who are found to use tactile/optical cues to know the presence of an epibiont, the snails do it through chemical cues. All the snails in our experiment latched on to the prey and continued with the predation. We did not encounter any event where the snail abandoned the prey after initial inspection. This implies that the barnacles make it difficult for predators to find a suitable drilling site. The position of the attachment seems to influence the handling time; the closure it is to the drilling site, the longer it takes to complete the drilling (Fig. 5A). The size of the encrustation, however, does not play any role (Fig. 5B). Our observations support an increase in handling time due to presence of encrusters and hence, the avoidance can be explained by the optimal foraging theory.

### Evolutionary implications

Predation by drilling gastropods has been claimed to be an important selective force shaping the evolutionary trajectory of the marine benthos, especially of barnacles [44]. A significant reduction in the number of wall plates in Balanomorph barnacles has been attributed to the post-Cenozoic rise in drilling predation [Fig. 5 in 44]. Fossil record of predation, however, shows a low drilling frequency of barnacles (DF=0.1) [Fig. 2 in 39] in contrast to the heavily targeted molluscan prey (DF=0.3) [Fig. 2 in 3]. Such low predation intensity may also reflect the non-overlapping nature of habitat shared between the common drilling predators and the barnacle prey. Majority of the drillings on barnacles are created by muricid drillers. Naticids, a very common driller of molluscs, have rarely been reported to drill modern barnacles in a rocky substrate, probably because naticids live mainly infaunally [40, but see 45], whereas barnacles are epifaunal encrusters. In contrast, the barnacles have been found to be more likely to be drilled when they are in association with a mobile molluscan prey [37]. In spite of the rarity of the naticid predation on barnacles, we cannot ignore the potential role of naticid predation on the evolution of the barnacle-mollusc association, especially the association with the epifaunal/semi-infaunal molluscan prey of intertidal habitat where the naticid predators are abundant [37, 38]. Barnacle infested molluscan prey enjoys an associational resistance against drilling predation, probably at the cost of an increased mortality of barnacles. The increased fitness of the barnacle infested individuals of prey species would likely to favor the survival of individuals with suitable morphological features conducive to attract epibionts.

Epibiosis on a shelled organism depends on a number of factors including host size and shell topography. A study on extant brachiopods suggests that epibionts prefer to colonize species with pronounced shell ornamentation over the smooth shelled species [46]. Vance [10] also showed that clams with intact concentric lamellae had significantly more epibionts on its shell than smooth bivalve of the same species. Our results demonstrate that an association with epibionts like barnacles may offer effective resistance against drilling predation. It is known that predation often triggered an evolutionary trend of highly ornate shell architecture [47]. Apart from the direct benefit of resisting predation induced damage, such evolutionary trend of higher sculpting may also be influenced by its ability to attract larger colony of epibionts. This would especially be true for immobile epifaunal whose inherent lack of mobility is not affected by the extra shell mass of the epibionts. Because highly sculpted shells are more suitable for encrustation [10, 46], sculptures may even be exapted in molluscan prey to facilitate epibiotic attachment given their effectiveness in deterring drilling predation on molluscs. Such exaptation of shell ornamentation is not uncommon. Some ornamental features such as concentric ribs have been claimed to be an exaptation against drilling predation (in contrast to an adaptation) because of their pre-Cretaceous existence well before the Cenozoic rise of drilling predation [48]. It requires a thorough investigation to compare of predation intensity of encrusted prey in the fossil record and their shell ornamentation to resolve this issue.

## Acknowledgements

We are grateful to Anupama C, Subhasmita Swain, Deepjay Sarkar, Madhura Bhattacherjee and C Venkateswar Reddy for their help in specimen acquisition from the field and in setting up the experiment. DC acknowledges the financial support of SERB (CRG/2018/002604).

